# GenSLMs: Genome-scale language models reveal SARS-CoV-2 evolutionary dynamics

**DOI:** 10.1101/2022.10.10.511571

**Authors:** Maxim Zvyagin, Alexander Brace, Kyle Hippe, Yuntian Deng, Bin Zhang, Cindy Orozco Bohorquez, Austin Clyde, Bharat Kale, Danilo Perez-Rivera, Heng Ma, Carla M. Mann, Michael Irvin, J. Gregory Pauloski, Logan Ward, Valerie Hayot-Sasson, Murali Emani, Sam Foreman, Zhen Xie, Diangen Lin, Maulik Shukla, Weili Nie, Josh Romero, Christian Dallago, Arash Vahdat, Chaowei Xiao, Thomas Gibbs, Ian Foster, James J. Davis, Michael E. Papka, Thomas Brettin, Rick Stevens, Anima Anandkumar, Venkatram Vishwanath, Arvind Ramanathan

**Author notes:** Joint first authors (Maxim Zvyagin, Alexander Brace, Kyle Hippe, Yuntian Deng). Permission to make digital or hard copies of all or part of this work for personal or classroom use is granted without fee provided that copies are not made or distributed for profit or commercial advantage and that copies bear this notice and the full citation on the first page. Copyrights for components of this work owned by others than ACM must be honored. Abstracting with credit is permitted. To copy otherwise, or republish, to post on servers or to redistribute to lists, requires prior specific permission and/or a fee. Request permissions from. Supercomputing ‘22, November 14-19, 2022, Dallas, TX © 2020 Association for Computing Machinery. ACM ISBN ISBN.$15.00 https://doi.org/finalDOI.

## Abstract

We seek to transform how new and emergent variants of pandemic-causing viruses, specifically SARS-CoV-2, are identified and classified. By adapting large language models (LLMs) for genomic data, we build genome-scale language models (GenSLMs) which can learn the evolutionary landscape of SARS-CoV-2 genomes. By pretraining on over 110 million prokaryotic gene sequences and finetuning a SARS-CoV-2-specific model on 1.5 million genomes, we show that GenSLMs can accurately and rapidly identify variants of concern. Thus, to our knowledge, GenSLMs represents one of the first whole genome scale foundation models which can generalize to other prediction tasks. We demonstrate scaling of GenSLMs on GPU-based supercomputers and AI-hardware accelerators utilizing 1.63 Zettaflops in training runs with a sustained performance of 121 PFLOPS in mixed precision and peak of 850 PFLOPS. We present initial scientific insights from examining GenSLMs in tracking evolutionary dynamics of SARS-CoV-2, paving the path to realizing this on large biological data.

## 1 JUSTIFICATION

We demonstrate using >1.63 Zettaflops, sustained performance of 121 PFLOPS (mixed precision) and 850 PFLOPS peak, in training one of the largest foundation models on whole genome sequences to characterize SARS-CoV-2 variants of concern. Our models will inform timely public health intervention strategies and downstream vaccine development for emerging variants.

## 2 PERFORMANCE ATTRIBUTES

| Performance Attribute | Our Submission |
| --- | --- |
| Category of achievement | Scalability; time-to-solution |
| Type of method used | Explicit; deep learning |
| Results reported on basis of | Whole application including I/O |
| Precision reported | Mixed Precision |
| System scale | Measured on full system |
| Measurement mechanism | Hardware performance counters; application timers; performance modeling |

## 3 OVERVIEW OF THE PROBLEM

Tracking of novel and emergent variants for viruses such as severe acute respiratory syndrome coronavirus 2 (SARS-CoV-2) has been enabled by rapid sequencing and sharing of whole genome sequence data (Otto et al., 2021). As of September 2022, >13 million SARS-CoV-2 genomes have been deposited in the GISAID repository^1^. SARS-CoV-2 represents one of the most deeply sequenced viral genomes and is therefore a rich source of information for understanding various factors that drive its evolution. Despite its slow mutation rate, over the past three years SARS-CoV-2 has evolved several variant strains containing unique mutation patterns, many of which lead to novel viral phenotypes including higher antigenicity, transmissibility, and fitness (Cosar et al., 2022).

This has prompted the US Centers for Disease Control and Prevention (CDC) to identify four SARS-CoV-2 variant categories, including: variants being monitored (VBM), variants of interest (VOI), variants of concern (VOC), and variants of high consequence (VOHC). Classification stems from SARS-CoV-2 growth dynamics and threat to pre-existing immunity (Cosar et al., 2022, Otto et al., 2021). Today, SARS-CoV-2 VOCs include B.1.1.7 (Alpha), B.1.617.2 (Delta), and B.1.1.529/BA.1-BA.5 (Omicron). Although deep sequencing of viral genomes across patient populations has enabled substantial progress, identifying variants is still tedious and resource-intensive, requiring costly laboratory tests and diagnostics. Together, these factors contribute to significant time expenditure to recognize and subsequently make informed decisions for public health intervention strategies (Baker et al., 2021).

Artificial intelligence (AI) and machine learning (ML) approaches promise to transform real-time pandemic monitoring (Syrowatka et al., 2021). Instead of reacting after the emergence of variants to identify VOCs over potentially several weeks (see Sec. 4.1), AI/ML techniques can leverage deep sequencing data to proactively identify mutations in viral proteins and characterize evolutionary patterns that can assist in predicting and describing future VOCs (Beguir et al., 2022, Hie et al., 2021). However, obtaining high-quality, global-scale genome datasets can be challenging, as diverse sequencing technologies can result in variable quality and coverage of sequenced genomes. Sequence-based feature extraction techniques followed by traditional ML approaches have demonstrated promise in the early identification of VOCs (Beguir et al., 2022, Maher et al., 2022, Wallace et al., 2022); however, they remain limited to sequence signatures of regions of interest in the genome. This unmet challenge presents an opportunity for effective whole genome-scale surveillance of global pandemics and early identification of VOCs, with the goal of enabling the development of robust public health intervention strategies prior to surges in case numbers and improving vaccine-design strategies on emerging variants.

We posit that by leveraging the recent success of large-language models (LLMs) in natural language processing (NLP) tasks (Wei et al., 2022), we can develop global-scale, whole genome surveillance tools. In this paper, we use LLMs to characterize SARS-CoV-2 evolutionary dynamics and reconstruct SARS-CoV-2 variant emergence. We adapt LLMs developed for understanding human languages to genomic sequences, called *genome-scale language models (GenSLM)*, and validate this approach in modeling VOC assignments for SARS-CoV-2 using historical data. Our contributions include:

- We develop the largest *biological* LLMs with codon tokenization (with 2.5 and 25 billion trainable parameters) to date, trained across a diverse set of 110 million prokaryotic gene sequences. These are the first foundation models trained on raw nucleotide sequences to demonstrate substantial improvement in predictive performance in identifying VOCs. We make these models and weights openly available to the scientific community ^2^.
- We design and validate a novel hierarchical transformer-based model that uses both Generative Pre-trained Transformers (GPT) (on individual gene sequences) and stable diffusion to capture the correct context and longer-range interactions in genome-scale datasets. This model enables us to prospectively model SARS-CoV-2 evolution by leveraging its generative capabilities.
- We showcase training foundation models on both conventional (GPU-based) systems (Polaris at ALCF and Selene at NVIDIA) and on emerging AI-accelerator hardware (interconnected Cerebras CS-2 systems), and demonstrate high watermarks for time-to-solution (model performance described by its perplexity or accuracy). In addition, we present scaling benchmarks, which demonstrate that training GenSLMs can be intensive—performing over 1.63 × 10^21^ floating point operations (a mix of FP16 and FP32; 1.63 Zettaflops) with a sustained performance of 121 PFLOPS (mixed precision) and 850 PFLOPS peak, over the course of training runs.

Together, these capabilities go beyond state-of-the-art techniques for global-scale whole genome surveillance of pandemic-causing viruses and address a critical infrastructure need for global public health organizations.

## 4 CURRENT STATE OF THE ART

Current approaches for tracking viral evolution rely on infectious disease specialists who examine variations, identify epitopes of interest (i.e., portions of the virus that elicit immune response), classify variants, and eventually flag them for further laboratory testing and analysis (Brouwer et al., 2020, Greaney et al., 2021, Ju et al., 2020, Zost et al., 2020). This process is widely used for tracking viral infections, including seasonal influenza (Doud et al., 2018). Identifying strains of interest helps prioritize downstream vaccine development workflows. However, this process is time-consuming and laborious. While data sharing in the community has enabled unprecedented progress in developing vaccines for pandemics such as COVID-19, there still exists an unmet challenge in accelerating the detection and prediction of viral VOCs via computational and experimental toolkits.

### 4.1 Early warning systems for viral evolution

Several early warning systems for tracking COVID-19 have been developed; however, they utilize case counts, internet search parameters, and other allied data focused on monitoring case counts in a local geographic area (Ramchandani et al., 2020). The Bacterial and Viral Bioinformatics Resource Center (BV-BRC)^3^ provides the SARS-CoV-2 Emerging Variant Tracking and Early Warning System, which enables users to browse current and past variant lineages and track their prevalence by isolation date, geographic location, and other metadata fields. A heuristic is used to compute month-over-month growth rates and highlight rapidly growing variants that may cause future infection surges. Mutations from each variant are mapped to known epitope sites and regions of the genome known to be involved in antibody escape to enable further assessment of mutation impact. Recently, Hie et al. (Hie et al., 2021) used protein language models (PLMs) and adapted concepts from NLP to model escape variants across three different viruses, including SARS-CoV-2. In each virus, they identified a certain protein (e.g., SARS-CoV-2 Spike/S protein) and modeled its evolutionary history using transformers to describe differences between ordinary variants and VOCs. Similarly, Beguir et al. (Beguir et al., 2022) leveraged a PLM to accurately classify VOCs; using experimental assays, they also validated these VOCs and demonstrated the ability to flag them in advance of World Health Organisation designation. However, viral evolution is not isolated to one protein but occurs at the genome scale. We propose a system that *learns to model whole-genome evolution patterns using LLMs based on observed data*.

### 4.2 Large language models (LLMs)

The introduction of transformers (Vaswani et al., 2017) — and subsequent LLMs such as Bidirectional Encoder Representations from Transformers (BERT) (Devlin et al., 2018) and generative pre-trained transformers (GPT) (Radford et al., 2018) — has revolutionized natural language understanding. These models have been used to generate text, speech (Gulati et al., 2020), and images (Han et al., 2022). They have also been employed to understand the language of nucleic acids (DNA/RNA) and proteins. Protein language models (PLMs), using amino acid alphabets, are the most heavily investigated biological LLMs (Elnaggar et al., 2022, Rives et al., 2021), with demonstrated success in downstream tasks such as protein function prediction (Unsal et al., 2022) and engineering (Ferruz et al., 2022). Nucleotide LLMs, using DNA/RNA alphabets, are still understudied (Avsec et al., 2021). Compared to the rich alphabet and short length of information-dense protein sequences that traditional attention models from NLP can successfully learn, nucleotide LLMs rely on much simpler alphabets and extremely long-range signal (e.g., across open reading frames or co-evolutionary patterns) and require significant domain adaptation to yield good results. When applied on the scale of entire genomes, GenSLMs also operate on much larger sequence lengths than are traditionally seen in NLP applications—the max sequence length for SARS CoV2 tasks was 10,240 tokens in comparison to the standard 1,024 or 2,048. Further, viral genomes often undergo frameshift mutations leading to differential translation, introducing ambiguity not present at the protein scale. Our work addresses these challenges by leveraging a hierarchical LLM: a generative pre-trained transformer to capture shorter/local (codon-level) interactions, and a diffusion-based model to capture longer-range interactions to describe the biological complexity of viral evolution (Sec. 5.2).

### 4.3 Workflow infrastructure

Scientific applications for HPC are increasingly written as a composition of many interconnected components. Application components may have different hardware or software requirements, run durations and execution frequencies, or dependencies with other components. Workflow systems such as Swift, Parsl, Balsam, and RADICAL Cybertools support the design of applications as directed graphs of tasks and manage their execution on available resources.

There is significant diversity in workflow implementation; e.g., Swift/T expresses workflows in bespoke programming languages that are compiled into an MPI program (Wozniak et al., 2013). Parsl, in contrast, is built on Python’s native concurrency library and dynamically constructs a task graph as a Python program is interpreted (Babuji et al., 2019). Balsam (Salim et al., 2019) and RADICAL CyberTools (Balasubramanian et al., 2019) rely on a central task database from which the launcher, running on compute resources, pulls and executes tasks. A centralized database enables state persistence across runs, and task dependencies can be defined as a DAG. Most workflow systems support interfacing with HPC job schedulers or cloud providers to acquire resources and transmit files between remote resources–key features for our use case.

Dynamic workflows, where new tasks are continually added in response to new results, are emerging as an extension of workflow managers. Many dynamic workflow systems, such as DeepHyper (Balaprakash et al., 2018) and RocketSled (Dunn et al., 2019), are purpose-built to solve optimization problems. LibEnsemble (Hudson et al., 2022) provides a more general interface where users decouple a dynamic ensemble into a “generator” which spawns new tasks based on results from a “simulator.” Toolkits such as Ray (Moritz et al., 2018) and Colmena (Ward et al., 2021) provide more flexible approaches where a number of “agents” can cooperatively coordinate tasks. These libraries handle *where* and *how* tasks are executed and provide useful abstractions so users can focus on component/task logic (i.e., *what* and *when)*.

## 5 INNOVATIONS REALIZED

Given the limitations of current approaches in identifying VOCs, there is a need to develop an integrated system that can automatically ‘learn’ features within the SARS-CoV-2 genome that distinguish VOCs, while also being able to generate new sequences that characterize emerging variants of the virus. We posit that by leveraging existing sequencing data on the virus, we can train LLMs that can model the SARS-CoV-2 evolutionary trajectory. Training LLMs on SARS-CoV-2 genome datasets is non-trivial due to the need to: (1) address the limitations of training LLMs with genomic sequences; and (2) overcome infrastructural challenges to enable LLM training on large sequence lengths in a reasonable time.

### 5.1 Data collection and description

#### 5.1.1 SARS-CoV-2 genome dataset

The Bacterial and Viral Bioinformatics Resource Center (BV-BRC) web resource provides integrated data and analysis tools for bacterial and viral pathogens to support infectious disease research. It hosts >600,000 bacterial genomes and 8.7 million viral genomes, including 6.4 million SARS-CoV-2 genomes. All SARS-CoV-2 genome sequences were acquired from NCBI’s GenBank and SRA databases and uniformly annotated using VIGOR4 (Wang et al., 2012) to provide accurate and consistent annotation of open reading frames (ORFs) and mature peptides across all SARS-CoV-2 genomes. Automated and manual curation provided accurate and uniform metadata across all genomes, including host name, geographic location, and collection date.

To build GenSLMs for detecting and predicting SARS-CoV-2 variants of interest, we used >1.5 million high-quality BV-BRC SARS-CoV-2 complete genome sequences. We filtered out any genome sequences with < 29,000 bp and >1% ambiguous bases. However, we note here that the data collected might not have sufficient diversity— meaning that any model trained on the SARS-CoV-2 dataset may end up overfitting to the data, with little opportunity to generalize. Hence, we took a foundation model-based approach that allowed us to first build a more general model using a much larger collection of diverse genomic data, namely gene-level data from prokaryotes.

We also utilized a dataset collected by the Houston Methodist Hospital System - one of the largest single-institution collections of SARS-CoV-2 genome sequences in the United States. We started here with 70,000 SARS-CoV-2 patient samples collected from May 15, 2020 to January 14, 2022 and sequenced on Illumina instruments. To ensure high quality, we first masked the leading and trailing 100 nucleotides for each sequence, as well as 56 positions in the spike protein-encoding region that had low depth due to poor primer binding. Sequences with >256 ambiguous characters were discarded, leaving 16,545 total sequences. This subset was used for building phylogenetic analyses at genome-scale (see Sec. 5.3).

#### 5.1.2 BV-BRC dataset

To allow for better generalization and to avoid overfitting of models to the SARS-CoV-2 data, we used >110 million unique prokaryotic gene sequences from BV-BRC. The BV-BRC database provides cross-genera protein families, PGfams, which allow for collection of homologous gene or protein sequences across taxa that perform the same biological function (Davis et al., 2016). We queried BV-BRC to find 10,206 unique PGfams, each with >30,000 unique members. For each PGfam, we collected high-quality non-redundant gene and protein sequences after filtering out sequences that were more than one standard deviation from the PGfam’s mean length. We term the Genome-Scale Language Models (GenSLMs) models trained on this data *foundation models*.

### 5.2 Large language models

Large-language model (LLM) training required both algorithmic and performance-level innovations. For algorithmic innovations, we describe two key limitations of current LLMs. For performance innovations in achieving optimal time-to-solution (training time to achieve some accuracy or perplexity), we leverage an interconnected cluster of four Cerebras CS-2 AI accelerators and scale to large GPU-based supercomputers to train our LLMs.

#### 5.2.1 Genome-scale Language Models (GenSLMs)

We introduce GenSLMs as a means to go beyond current PLMs to describe evolutionary dynamics of SARS-CoV-2. Instead of focusing on specific proteins, GenSLMs leverage genome-scale data to model individual mutations at the nucleotide scale, thus implicitly accounting for protein-level mutations at the codon level. Fig. 1 shows that GenSLMs take nucleotide sequences of SARS-CoV-2 genomes as input and learns a semantic embedding of individual codons, which can then be *translated* to the 29 individual protein sequences that are encoded by the virus.

**Figure 1:**
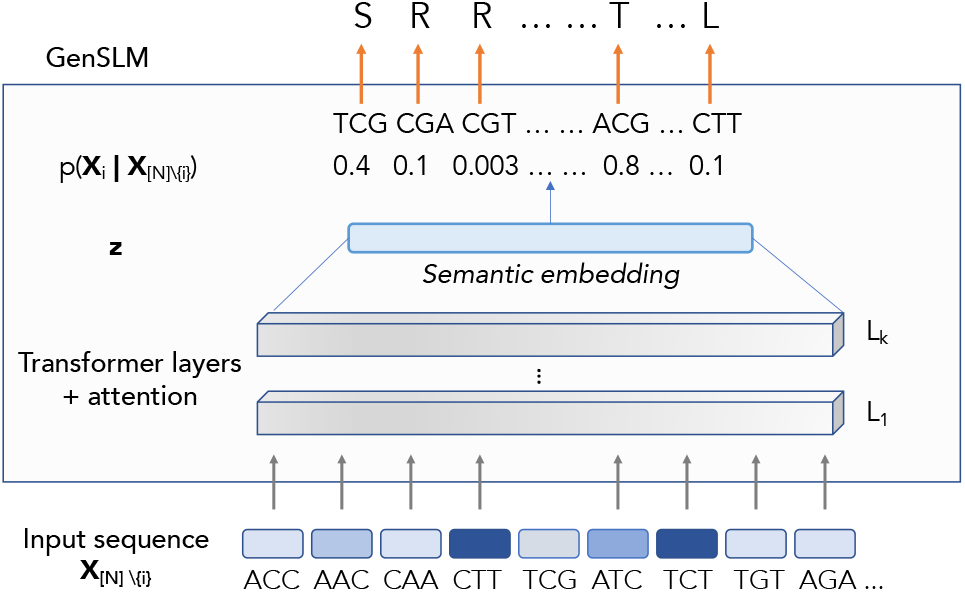
Overview of GenSLM models for predictive modeling of SARS-CoV-2 evolution. The inputs to GenSLM are nucleotide sequences, encoded at the codon level (every three nucleotide represents a codon; hence the 20 natural amino acid language is described by 64 codons). These inputs are successively fed into transformer blocks (referred to as layers (*L_i_*)), which ultimately results in learning a semantic embedding 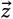 space from which one may obtain the probability of any given sequence token 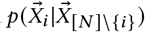, where *N* represents the sequence length and *i* represents a particular position in the entire genome.

However, there are two fundamental challenges when training GenSLMs directly from SARS-CoV-2 genome sequences: (1) The entire genome consists of ~30,000 nucleotides (which translates to ~10,000 codons/amino-acids). LLM training on long sequences can be challenging because attention mechanisms largely focus on shorter/local segments of the genome rather than global patterns. (2) The overall sequence similarity in SARS-CoV-2 genomes is high (>~99%), with only a small (yet significant) number of changes that yield distinct phenotypes. Thus, there is a need to address diversity in the sequences such that the trained model can generalize. It is also necessary to account for frameshifts in viral genomes.

To overcome these challenges, GenSLM implicitly recognizes intrinsic hierarchy (based on the central dogma) of individual protein production via DNA transcription and mRNA translation. We trained on gene-level data from BV-BRC (see Sec. 5.1) to mimic this process with GenSLMs. Although mapping between codons and amino acids is degenerate (multiple codons may encode the same amino acid) (Shin et al., 2015), we posited that with sufficient diversity in the dataset, GenSLMs could exploit intrinsic organization of gene-level data to learn biologically-meaningful latent representations. The training process follows a procedure similar to the one outlined in (Zhang et al., 2022). We refer to the models trained on the BV-BRC dataset as GenSLM foundation models.

While the benefits of pre-training LLMs on natural text are well known (Turc et al., 2019), obtaining the optimal number of transformer layers and training on such a diverse set of gene data were challenging. We, therefore, trained GenSLM foundation models on a wide set of parameter scales ranging from 25 million to 25 billion, with a maximum sequence length of 2,048 tokens on the BV-BRC dataset. Additionally, to evaluate performance on downstream tasks, we fine-tuned the foundation model GenSLMs using a maximum sequence length of 10,240 tokens for the 25M and 250M model sizes on the SARS-CoV-2 datasets (see Table 1). We note that for the larger model sizes (2.5B and 25B), training on the 10,240 length SARS-CoV-2 data was infeasible on GPU clusters due to out-of-memory errors during attention computation.

**Table 1:** Description of GenSLMs foundation model architectures. #H – number of attention heads; #L – number of layers; *d*_model_ – embedding size; LR – learning rate (if range is specified, decayed by factor of 10 each update); B – global batch size in number sequences per step; P – total number of trainable parameters; MSL – maximum sequence length. The ^†^ denotes models that we were also able to train on the 10,240 sequence length for the full genome.

| | #H | #L | $d_{\text{model}}$ | LR | B | P | MSL |
| --- | --- | --- | --- | --- | --- | --- | --- |
| GPU | 8 | 8 | 512 | $5e^{-05}$ | 4096 | 25M | 2048$^\dagger$ |
| | 16 | 12 | 1,840 | $5e^{-05}$ | 4096 | 250M | 2048$^\dagger$ |
| | 64 | 26 | 3,968 | $5e^{-05}$ | 512 | 2.5B | 2048 |
| | 64 | 64 | 8,192 | $5e^{-05} - 5e^{-09}$ | 1024 | 25B | 2048 |
| | #H | #L | $d_{\text{model}}$ | LR | B | P | MSL |
| CS-2 | 12 | 12 | 768 | $2.8e^{-04}$ | 33 | 123M | 10240 |
| | 12 | 12 | 768 | $2.8e^{-04}$ | 132 | 123M | 10240 |
| | 16 | 24 | 2048 | $7.5e^{-05}$ | 11 | 1.3B | 10240 |
| | 16 | 24 | 2048 | $1.5e^{-04}$ | 44 | 1.3B | 10240 |

The entire repertoire of results from the GenSLM foundation models is beyond the scope of this paper. However, as an empirical demonstration of the power of GenSLMs trained on the SARS-CoV-2 genomes, the learned latent space projected onto a lowdimensional manifold as determined by t-distributed Stochastic Neighbor Embedding (t-SNE) meaningfully distinguishes the SARS-CoV-2 variants as shown in Fig. 2. This observation is significant because this GenSLM-25M model was specifically trained only on the first year of the SARS-CoV-2 data (consisting of ~ 85,000 SARS-CoV-2 genome sequences) – meaning that the model did not have the opportunity to see any of the other strains. Thus, the ability of GenSLM to generalize and distinguish between SARS-CoV-2 variants implies that the learning process is robust and the underlying features can generalize to downstream tasks. We also note that as the model parameters increase, the perplexity of the model also improves, agreeing with previous observations (Radford et al., 2018).

**Figure 2:**
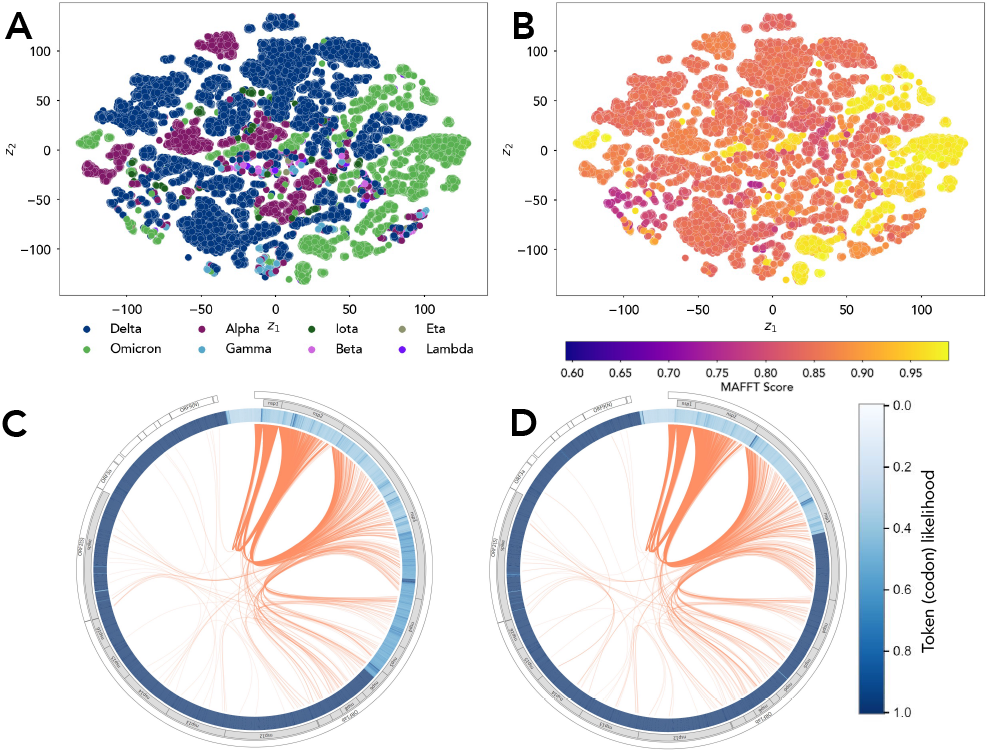
GenSLMs learned latent space describes biologically meaningful properties for SARS-CoV-2 genomes. (A) The embeddings from GenSLMs are visualized with t-distributed stochastic neighbor embedding (t-SNE) and each gene sequence is represented as a dot in the 2D plot. We paint each sequence by its variant ID – although we have more than 515 PANGO (Rambaut et al., 2020) lineages represented in the data, we only show those with WHO designated labels. (B) The latent space can also be painted with the MAFFT-determined alignment score (Yamada et al., 2016) with respect to an Omicron genome; clustering in the distance measures is clearly visible. Visualizing the sequence log-likelihood (blue bar) and the cross-protein attention (orange lines) from (C) Delta and (D) Omicron SARS-CoV-2 strains highlights how different the co-evolutionary patterns are in these lineages. It is interesting to note that while the Spike protein from Delta strain shows coupling to nsp3, nsp5, and other proteins, these couplings are not observed in the Omicron strain.

We note however that these training runs frequently take >1 week on dedicated GPU resources (such as Polaris@ALCF). To enable training of the larger models on the full sequence length (10,240 tokens), we leveraged AI-hardware accelerators such as Cerebras CS-2, both in a stand-alone mode and as an inter-connected cluster, and obtained GenSLMs that converge in less than a day (Sec. 5.2.4).

#### 5.2.2 Reward-guided beam search for generative modeling

A subsequent use of the GenSLM models is in its ability to generate new SARS-CoV-2 sequences, with the eventual goal of predicting yet unseen VOCs. One challenge with such sequence-based generation strategies is sampling sequences with particular properties. Given a conditional sequence model *p_θ_* with weights, *θ*, the most likely sequence is 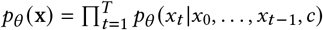 where *c* is the context from the previous inference. However, computing this directly is generally intractable as it is 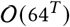, where *T* is the maximum sequence length with a vocabulary of size 64. Heuristics like greedy sampling are commonly used, where a sequence is generated iteratively, with the next token *x_t_*, maximizing *pθ*(*x_t_*|*x_0_*,…,*x*_*t*-1_, *c*) with complexity 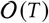.

Beam search is standard practice, combining a search strategy with a heuristic where *k* is the number of beams explored with complexity 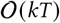. First, *k* samples are drawn with the highest probability (or sampled from a multinomial distribution) and added to the set of possible hits 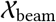. Let 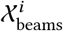 be the set of beams of length *i*. Then, for time step *t*, select *k* tokens *x_t_* from the set of all tokens which score highest (or sampled from multinominal distribution) via 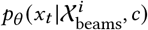. The highest scoring beams from 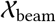 are selected via 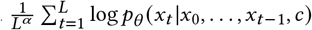 and output, where *L* is the length of a sequence and *α*, is a length penalty.

Given an episodic reward function 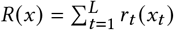, we modify the scoring function for beam search with

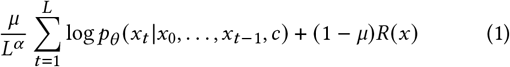

where 0 ≤ *μ* ≤ 1 is a hyperparameter. Since the reward function is episodic, at each step of beam search, the highest scoring beams are chosen with

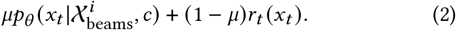

This scoring modification effectively alters the likelihood of tokens to be sampled based on maximizing the reward function. In order to sample sequences that are similar to a fixed sequence *y*, we utilize *r_t_*(*x_t_*) equal to the global alignment score between *y_t_* and *x_t_* (Needleman and Wunsch, 1970). This scoring bias modification effectively implements a property scoring function into beam search without altering the complexity of beam search sampling. In the case of non-episodic reward functions, rewards can only be computed at the final time step in eq. 1.

#### 5.2.3 Diffusion-based hierarchical modeling

Token-level autoregressive modeling has difficulty in generating coherent long sequences due to its underlying challenge in capturing long-range dependencies (Papalampidi et al., 2022, Sun et al., 2021, 2022). We developed a new hierarchical-modeling method based on a latent-space diffusion model that operates on the ‘sentence’ level. For each genome sequence, we uniformly truncated every 512 codons by a special separator symbol; these 512 codons are considered a ‘sentence.’ (We set 512 codons to be a sentence such that the average number of sentences per sequence, around 20, matches the number of ORFs and non-coding regions: around 17.)

Our diffusion-based hierarchical modeling method consists of three parts: **(1) Learning high-level representations:** We trained a new encoder to embed sentences into a latent space with a contrastive loss such that learned features better capture high-level dynamics. Our contrastive loss is similar to the masked language modeling objective used in SpanBERT (Joshi et al., 2020), where we predict missing sentences in the middle by conditioning on the previous and the next sentences, and, at the same time, using randomly sampled sentences as negatives (distractors).

**(2) Modeling high-level dynamics with a diffusion model:** Given the encoder output of each genome, i.e., a sequence of sentence embeddings, we train a diffusion model to learn their distribution. The diffusion model parameterizes the distribution of high-level representations by applying a sequence of denoising operations on top of Gaussian noise. Similar to previous work (Ho et al., 2020, Vincent, 2011), we used denoising score matching as the training objective; we gradually apply noise to desired target representations and the diffusion model learns to denoise at each step.

**(3) Fine-tuning LMs with high-level planning:** Similar to Time Control LM (Wang et al., 2022), we fine-tuned GenSLMs as the decoder to generate the genome sequence conditioned on high-level representations learned in step (1), which we term the ‘high-level plan’. The decoder predicts the current codon token using previous codon tokens within the context window size and the corresponding sentence embeddings. The training objective is the same as in training the original GenSLMs.

The overall generation procedure is shown in Fig. 3. Note that without the guidance of the high-level representations z_0_, the decoder can only take into account a limited amount of context, but with the guidance of z_0_, the decoder can take into account long-term context because z_0_ is modeled globally.

**Figure 3:**
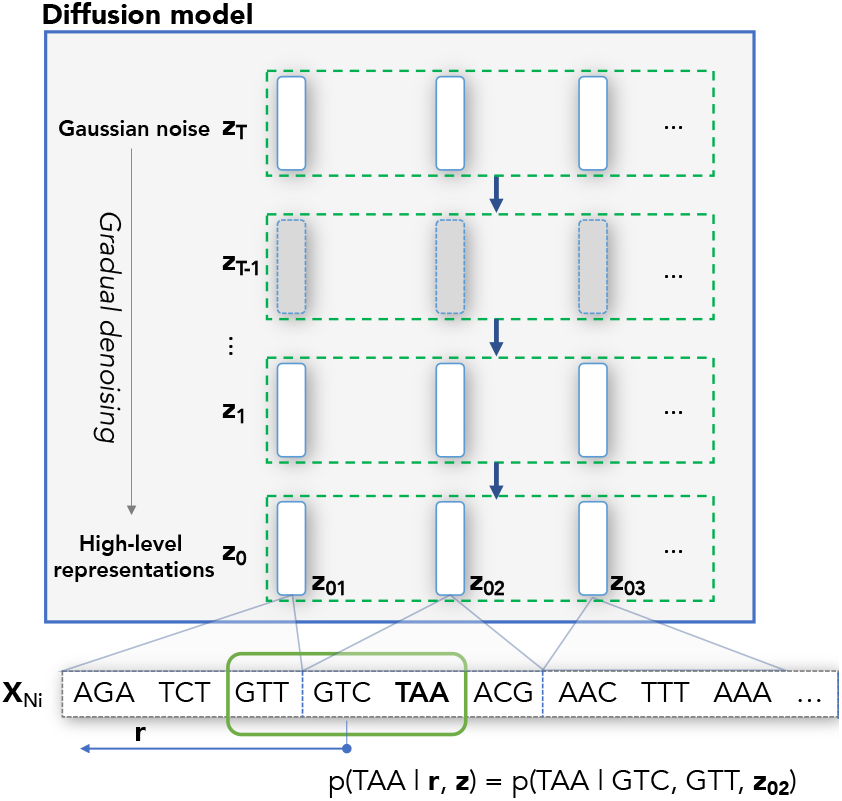
Illustration of diffusion-based hierarchical modeling. To predict a codon (such as TAA), we use both the previous codons within the context window (we use size 3 shown in green for illustration) and the high-level representations z.

We conducted experiments by training a baseline LM and a diffusion-based hierarchical LM on the 1.5M SARS-CoV-2 genomes (see Sec. 5.1.1). The goal of this experiment was to primarily assess if the diffusion model can ‘stitch’ together the context of the genes together at the genome-scale (much like how words are ordered in a sentence). The baseline LM is the hierarchical LM without high-level guidance - essentially, a normal transformer language model. We initialized both the baseline LM and the hierarchical LM decoders from the 2.5B foundation model trained on individual genes from BV-BRC (see Sec. 5.1.2). We used a context window size of 1,024. The sentence encoder is initialized from the 25M foundation model. The diffusion denoising model is a transformer with the same architecture as BERT (Kenton and Toutanova, 2019). We used 10 nodes from Polaris@ALCF for training, with a total of 40 A100 GPUs. We used an Adam optimizer with a learning rate of 1e-4, a batch size of 2, and trained for 13k updates. Training took approximately 6 hours. At generation time, we used a sliding window-based approach: we first generate 1,023 codons from a start-of-sequence symbol, then move the window 512 codons to the right, generate the next 512 codons, and repeat this process until either end-of-sequence is generated or a maximum of 15k codons have been generated.

To evaluate if the generated samples capture high-level dynamics, we compared the distribution of the number of 5’-3’ ORFs on real data and on 1,000 samples from the model. As shown in Fig. 4A, the diffusion-based hierarchical model outperforms the baseline LM in generating realistic ORFs, possibly due to the high-level plan, whereas the baseline LM can only account for the previous 1,023 codons. We also display the phylogenetic tree (see Sec. 5.3) of the generated sequences from the diffusion-based hierarchical model against real genomes in Fig. 4B. The plot exhibits that the generated sequences cover the different lineages including all the variants. Note that sequences with >120 mutations (1.4% of all generated sequences) were excluded; this further demonstrates that the diffusion-based hierarchical model can generate sequences that are of higher quality than the standard transformer-based model.

**Figure 4:**
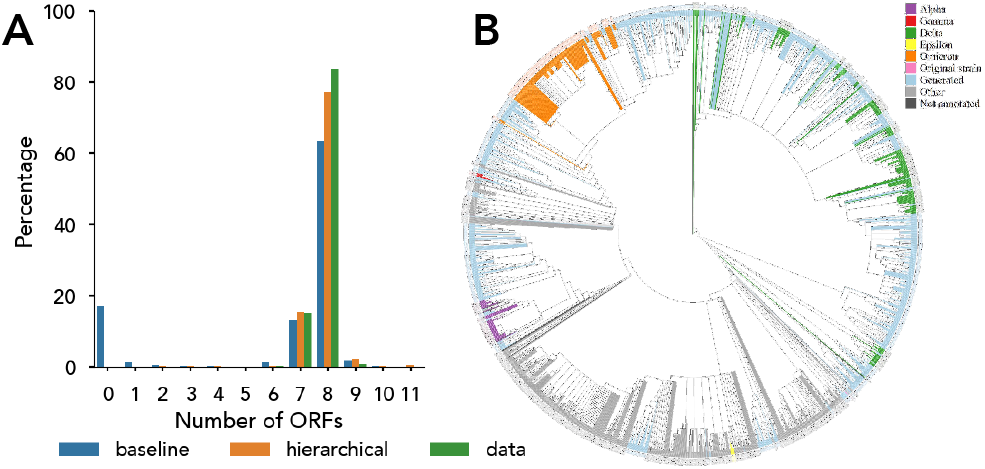
Diffusion-based hierarchical modeling of SARS-CoV-2 genomes results in generation of sequences that captures the correct context of various open reading frames (ORFs). (A) Comparison of statistics measured on generated sequences and on real data for the ORFs. Diffusion-based hierarchical LM has a global high-level plan whereas the baseline can only take into account the previous 1023 codons. (B) Generated sequences (light blue) from the model overlaid on the phylogenetic tree demonstrate that these sequences are similar to observed strains.

#### 5.2.4 Training with full viral genome sequences on Cerebras Wafer-Scale Cluster

Training LLMs on whole SARS-CoV-2 genomes with dense attention is challenging when using traditional approaches and hardware. With codon-based encoding, the model needs to handle sequences of 10,240 tokens. This results in high memory and computational demand, severe limitations to batch sizes to fit on a single device, and thus a need to develop and orchestrate complicated hybrid parallelism approaches to get reasonable performance with clusters of traditional devices. We overcome these challenges with the Cerebras Wafer-Scale Cluster (Hall et al., 2021), where it is possible to use only simple data parallelism, and achieve linear weak scaling, even when LLMs are trained on very long sequences. We pre-trained two GenSLMs to convergence on full viral genomes with dense attention (Table 1) using a sequence length of 10,240 codon tokens on a single CS-2, and on a cluster with four CS-2s, achieving desired accuracy and perplexity results in less than a day. Beyond compute performance, the Cerebras Wafer-Scale Cluster provides high usability through the appliance workflow, where users no longer need to handcraft different parallelism choices for their given hardware and only need to specify the number of CS-2s to start data-parallel training. This flexibility allows faster experiment iterations without compromising performance. Training GenSLMs with multiple CS-2s is pioneering work with the Cerebras Wafer-Scale Cluster, which demonstrates the potential of dedicated AI hardware to apply LLMs on long-range context and work with genome sequences at scale.

### 5.3 Phylogenetic analyses of whole genomes

As described in Sec. 5.1.1, we used a set of 16,545 sequences from the Houston Methodist Hospital System that were filtered for high-quality in order to analyze GenSLM outputs. We selected a diverse subset by embedding these sequences, tessellating the embedding space using a Gaussian mixture model (*N* = 40), and then sampling each tessellation using a uniform distribution, resulting in a set of 1,000 sequences maximizing coverage of the embedding space.

The 1,000 sequence subset was aligned to the NC_045512.1 severe acute respiratory syndrome coronavirus 2 isolate Wuhan-Hu-1 complete genome sequence using Mafft v7.310 (Yamada et al., 2016). We then generated a Newick-format phylogenetic tree from the alignment using RAxML Next Generation (Kozlov et al., 2019), which offers significant speed improvements over RAxML (Stamatakis, 2014). We then generated a phylogenetic tree using RAxML-NG’s “search” algorithm, which searches for a maximum-likelihood tree amongst 10 parsimonious trees and 10 randomly generated trees. This takes ~9 hours on 5 CPUs (the recommended RAxML-NG parallelization settings for our data.) We used the most likely tree generated as a seed tree for running further analyses with UShER.

UShER (Ultrafast Sample placement on Existing tRee, (Turakhia et al., 2021)) is a SARS-CoV-2-specific analysis tool that can quickly place new SARS-CoV-2 genomes onto an existing SARS-CoV-2 phylogenetic tree on the basis of mutation tracking. In addition to to a “seed” phylogenetic tree, UShER requires a variant call format (VCF) file to track mutation data, which we generated from our multiple sequence alignment by using snp-sites ((Page et al., [n.d.])). UShER stores the mutation information along with the tree in Google’s protobuff format created from the VCF and tree files.

We then used this tree as the basis for quickly examining and labeling generated sequences of interest. Generated sequences are (1) converted to fasta format, (2) aligned to the NC_045512 reference genome sequence, (3) mutation profiled using snp-sites, (4) placed on the seed phylogenetic tree using UShER, (5) proposed a variant label on the basis of the labels of its K nearest tree neighbors (where K=20 in our analyses), and (6) flagged for further examination if the sequence has the longest phylogenetic distance to the NC_045512 reference genome amongst its X nearest neighbors.

We chose this flagging scheme to select for sequences that were more distant to the original strain than the other sequences in their close lineage, as these sequences represent more heavily mutated novel genomes that may be more likely to produce variants of interest or concern.

### 5.4 Integrated visualization

When visualizing long-distance genomic relationships, a linear layout created edges crossing over entities and affected readability. We therefore developed a circular arrangement to visualize relationships between entities and positions. The visualization is flexible and supports a rich set of layers to encode various data properties. The outermost layer shows ORFs along the SARS-CoV-2 genome, while the next layer displays protein-coding locations. These interactive layers allow users to select an ORF or protein for closer examination. All other layers (except the innermost) display different properties of codons in the gene sequence. The layers encode codon properties and are presented with custom visualizations based on the property type; e.g., Fig. 2C and 2D, where probabilities of codons are encoded using a radial bar chart (intensity of the color represents the probability). The innermost layer visualizes the GenSLM attention relationships between codons. To reduce visual clutter, we employed hierarchical edge bundling techniques.

### 5.5 Workflow infrastructure

As illustrated in Fig. 5, we implemented and scaled the reward-guided beam search procedure (see Sec. 5.2.2), leveraging a workflow that couples (1) a GenSLM sequence generator, and (2) a Bayesian optimizer to tune the reward mixing hyperparameter *μ* to bias the generator towards a target property. On startup, an ensemble of GenSLM generators are submitted to perform an initial grid search over the *μ* ∈ (0,1) parameter space, providing sequences to update a Gaussian process surrogate model. This, in turn, suggests new *μ* values throughout the duration of the run. Parameters are chosen by random sampling of *n* points, and are scored by their negative expected improvement (the optimization is a minimization). Each generation task uses a single A100 GPU on Polaris to run an instance of the a 25M parameter GenSLM model, whereas the optimizer task uses the CPUs on a single node.

**Figure 5:**
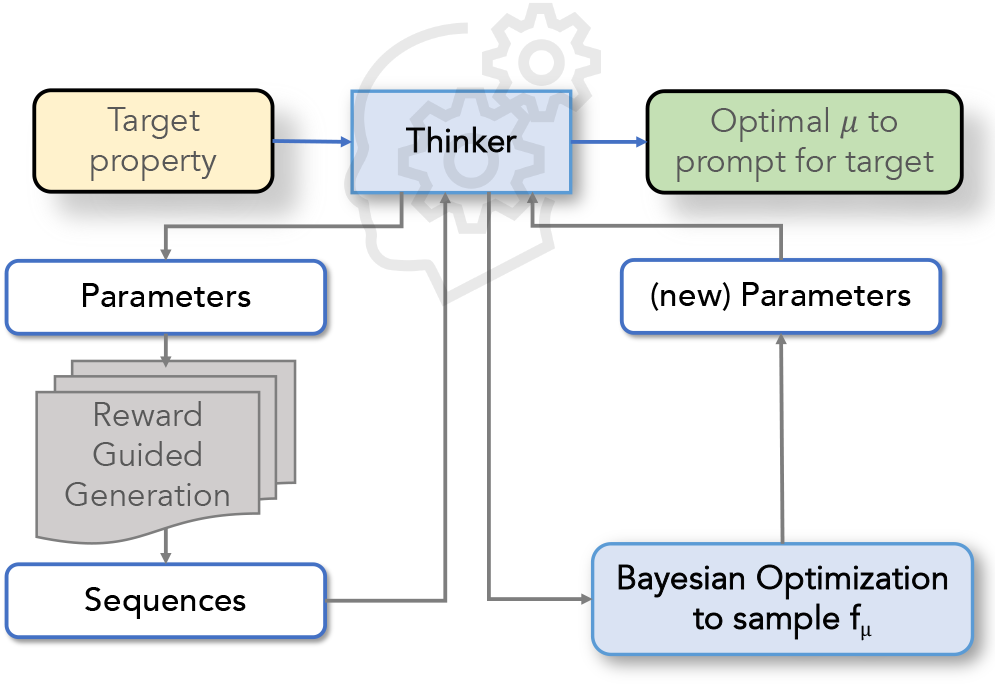
Conceptual overview of our workflow. A “Thinker” orchestrates data flow between two applications, namely the sequence generator and the Bayesian optimization to drive the generated sequences towards a target property using reward-guided beam search, where *μ* represents the mixing constant used to balance the reward function against the log likelihood of generating the next token.

We extend the Colmena workflow toolkit by implementing an Application abstraction for each workflow task (component). The Application provides for (1) inter-process communication when tasks are externally executable programs, and (2) warm-able functions to avoid duplicate initialization. The Application abstraction enables us to isolate the many generator instances from the Bayesian optimizer such that a single Thinker, executed on the login node, orchestrates communication and task submission to drive the property optimization. Leveraging Colmena allows us to implement concisely a multithreaded Thinker where one thread is responsible for handling outputs from the sequence generators and immediately submitting a new generation request to maximize utilization of the workers. This thread then handles any potential task failures by checking the return status and allows the workflow to be robust to application-level failures due to uncaught exceptions and hardware failures. The successful results are placed onto a queue where another thread reads and batches the results, (*μ*, sequence) pairs, for submission to the Bayesian optimizer application.

To further improve utilization of the workflow, we augment the Thinker with a inference-only copy of the surrogate model which is periodically transferred via pickling from the Bayesian optimizer application.

Workflows expressed with Colmena contain three components: a Thinker, a task server, and one or many workers. The Thinker defines the policies of the workflow, i.e., the dynamic dispatching of tasks and consumption of results. The Thinker is composed of agents; agents interact with each other and the task server via shared data structures. The task server pulls task definitions (task name and input pairs) from a task queue and executes tasks on workers via Parsl. The task server communicates task results from workers back to agents via a results queue.

For large task inputs or results, Colmena provides integration with ProxyStore (pro, 2021), a library for decoupling data movement from control flow. Task inputs or results that exceed a user-defined threshold are automatically communicated to the worker executing the task via more optimal means (e.g., file system or Redis server). This reduces overheads in the task server and workflow manager and enables lower latency task execution and higher throughput.

## 6 HOW PERFORMANCE WAS MEASURED

We evaluate the performance of GenSLM models on a diverse set of systems. We first explore the performance on two leadership class GPU-based supercomputing systems: 1) Polaris supercomputer at the Argonne Leadership Computing Facility (Polaris@ALCF), and 2) Selene supercomputer at NVIDIA (Selene@NVIDIA). Next, we evaluate the performance on the Cerebras CS-2 wafer-scale cluster.

In the June 2022 Top-500 list (Top500, 2022), Polaris is ranked at #14 with a peak of 44 PFLOPS and Selene is at #8 with a 63.4 PFLOPS peak. Table 2 compares the two systems used for evaluation. The Polaris system is an HPE Apollo Gen10+ system with 560 nodes interconnected with HPE Slingshot 10 using a Dragonfly topology. Each node consists of an AMD “Milan” processor with 32 cores with 512GB of system memory. Each node has four NVIDIA A100 GPUs—each with 40GB memory. Each node has two Slingshot-10 endpoints at 12.5 GB/s for the interconnect network. Selene is based on the NVIDIA DGX SuperPOD platform and consists of 560 nodes interconnected with Mellanox HDR fabric. Each node consists of two AMD “Rome” processors, each with 64 cores and 2TB system memory. Each node has eight NVIDIA A100 GPUs, each with 80GB memory. Each node has eight Mellanox ConnectX-6 HDR endpoints at 20 GB/s each for the interconnect network. Each A100 NVIDIA GPU is capable of achieving a peak of 19.5 TFLOPS in FP32, 156 TFLOPS in TF32, and 312 TFLOPS in FP16 and BF16.

**Table 2:** GPU supercomputing systems used for evaluation.

|  | Polaris | Selene |
| --- | --- | --- |
| June-2022 Top 500# | 14 | 8 |
| System size (nodes) | 560 | 560 |
| CPU | AMD Milan | AMD Rome |
| Sockets/Node (total cores) | 1 (32) | 2 (128) |
| System Memory (TB) | 0.5 | 2 |
| Number of GPUs per node | 4 | 8 |
| A100 GPU Memory (GB) | 40 | 80 |
| GPU Memory Technology | HBM2 | HBM2e |
| GPU Memory BW (TB/s) | 1.5 | 2.0 |
| Interconnect | HPE Slingshot-10 | Mellanox Infiniband HDR |
| NICs per node | 2 | 8 |
| Network BW per direction (GB/s) | 12.5 | 25 |
| Number of nodes (GPUs) scaled | 512 (2048) | 512 (4096) |

GenSLM was written with the PyTorch Lightning API (Pytorch, 2022), using transformer models from the Hugging Face repository (huggingface, 2022). PyTorch Lightning allows the use of several distributed training strategies to scale model training on clusters and supercomputers. This includes *DistributedDataParallel* and *DeepSpeed* (Rasley et al., 2020). We use mixed precision using FP16 and FP32 for our training runs. We focused our efforts on DeepSpeed, as its employment of various ZeRO strategies for optimization reduces the overall memory utilization in model training, particularly for large parameter models (Rajbhandari et al., 2020). Briefly, ZeRO strategies partition memory for training models— including the optimizer, gradient, and model states—to use aggregate memory across all GPUs. This enables training larger models on GPU-based systems and trades overall memory capacity for additional re-computation and communication. In particular, ZeRO-1 partitions optimizers across GPUs, ZeRO-2 partitions both the optimizers and gradients across all GPUs, and ZeRO-3 partitions the parameters, in addition to ZeRO-2 optimizations, across all GPUs. Additionally, ZeRO-3 can scale model sizes by leveraging CPU memory and any node-local storage to offload optimizer states, gradients, parameters, and optionally activations to CPU. We used PyTorch 1.12.0 and used NVIDIA NCCL 2.10.3 as the backend for DeepSpeed. We used an environment with Docker containers for the runs on Selene, and a bare-metal build using Conda on Polaris.

To measure compute performance of GenSLM model training, we use the DeepSpeed flops profiler (Deepspeed, 2022). The DeepSpeed flops profiler provides the flops and latency of the forward and backward passes and latency of the weight updates, and thus the compute performance of the GenSLM models. For scaling studies, we measure the entire end-to-end time including I/O as well as model training at scale. We measure achieved throughput in samples per second as the number of GPUs scales on the system. We use the NVIDIA Nsight tool (Bradley, 2012) to get an in-depth performance analysis.

### Cerebras Wafer-Scale Cluster

We also evaluated training performance on full viral genomic sequences on a Cerebras Wafer-Scale Cluster with four CS-2s (Hall et al., 2021). The Cerebras Wafer-Scale Cluster uses a weight streaming execution mode where weights are stored off-chip on MemoryX, a memory extension. Weights are streamed onto each CS-2 node using a broadcast/reduce fabric called SwarmX. Each CS-2 node is powered by the Wafer-Scale Engine, with 850,000 compute cores, 40 GB of on-chip memory, and 20 petabytes/s of memory bandwidth. After the computations, gradients are streamed back to MemoryX where weights are updated.

We used data-parallelism in the Cerebras Wafer-Scale Cluster through the appliance workflow, where no code changes or additional libraries were required to use either one or multiple CS-2 systems. GenSLM 123M and 1.3B were trained using the Cerebras reference implementation for GPT-2 model. This implementation is based on the TensorFlow estimator and is instrumented to collect accuracy, perplexity and throughput measurements. We worked with a Python virtual environment that included Cerebras software version 1.6. All training was done using mixed precision.

## 7 PERFORMANCE RESULTS

We evaluated the performance of scaling GenSLM training on the Selene and Polaris systems. We used two target sequence lengths (2,048 and 10,240) in our scaling studies. Fig. 6 depicts performance, in terms of overall throughput measured in samples/sec, as we scaled with the number of GPUs on both systems. In our runs, we used one rank per GPU with DeepSpeed ZeRO-3 optimizations. As we scaled the number of GPUs, we kept the batch size per GPU constant and scaled the global batch size appropriately. We modified the learning rate parameter to account for scaling the number of ranks. On Selene, for sequence length 2048, we employed twice the batch size used on Polaris for the 25M, 250M, and 2.5B models, as the A100 GPUs on Selene have twice the memory capacity compared to the Polaris GPUs. The performance obtained is the average of the throughput measured over multiple iterations.

**Figure 6:**
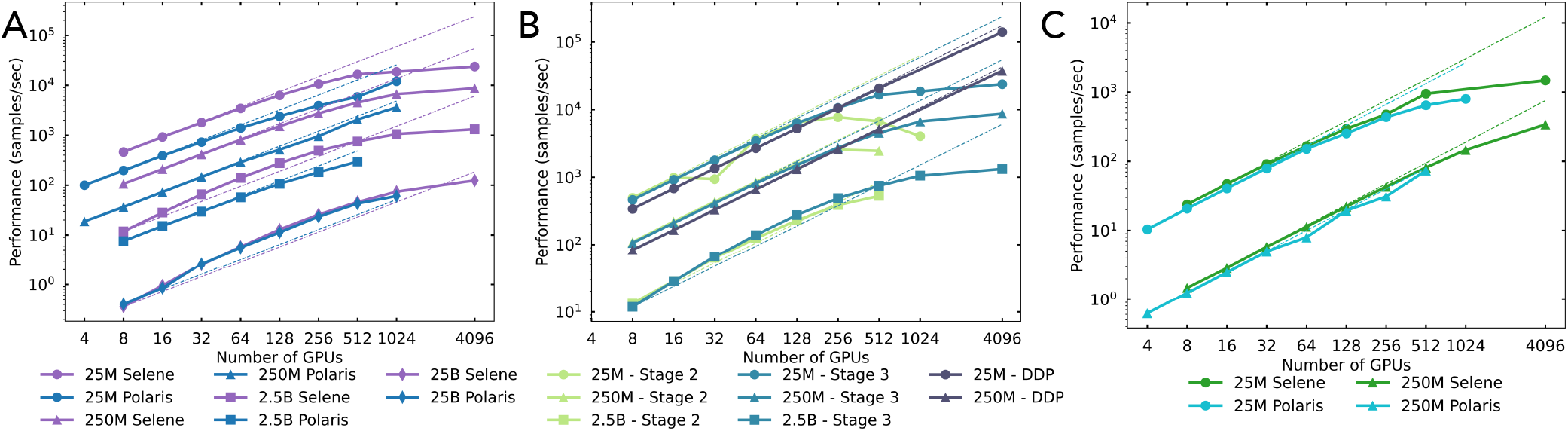
(A) Scaling results on Polaris and Selene systems for MSL=2048; (B) Scaling behavior of DDP vs. DeepSpeed runs on Selene (C) Scaling results on Polaris and Selene systems for MSL=10240;

We observed that as the model size increases from 25M to 25B, the total achievable throughput, in terms of samples/sec, decreases. This is expected as increasing the model size increases the computational, memory, and communication requirements. For the 25M, 250M, and 2.5B models, we observe a nearly 2× improvement in throughput on Selene in comparison to Polaris, as a double batch size is employed. In terms of efficiency, for smaller models, such as 25M, we observed a drop in scaling efficiency as we scaled beyond 256 GPUs. Two key attributes contributing to this include the fact that for smaller model sizes that run with ZeRO-3, the ratio of data movement to computational flops is too high to completely overlap these. We see better performance efficiency for larger models as they have higher utilization of computation and are able to better overlap communication with computation. Some inefficiencies here are also due to the performance of collectives and we investigate this further next. In the case of the 25B model, we are able to fit just a single batch on the GPU and observe a 50% improvement in the throughput achieved on Selene over Polaris. We attribute this to the increased interconnect performance on Selene together with the larger memory capacity. We observe a super-linear speedup for the 25B case on both systems as we scale to 1024 GPUs in comparison to the performance at 8 GPUs. This is attributed to the increased memory and data movement overheads at smaller GPU scales.

To gain detailed insights on the runs, we performed a profiling study on the 25M parameter model on both Polaris and Selene systems using the NVIDIA Nsight tool (NVIDIA, 2022). To account for the difference in the number of GPUs in a single node on both systems, we performed profiling runs with the 32 GPUs on 8 nodes on Polaris and 4 on Selene separately. We observed no significant delay between the steps/iterations—data loading and I/O were not bottlenecks. Given that the Selene DGX node has 80 GB memory compared with 40 GB on the Polaris node, it allowed doubling the batch size for the 25M parameter model, thereby achieving higher throughput than Polaris.

In addition, we performed a study comparing the scaling behavior of the distributed training framework implementations for PyTorch DistributedDataParallel (DDP) and with DeepSpeed with ZeRO Stage 2 and 3 on Selene. With DDP, we were constrained to smaller model sizes as it currently does not employ any memory optimization, unlike DeepSpeed. As seen from Fig. 6B, DDP-based runs exhibit linear behavior, while the performance of DeepSpeed runs saturated beyond 256 GPUs for the ZeRO-2 optimizer and 512 GPUs for the ZeRO-3 optimizations. For the 25M case, at 512 GPUs, DDP achieves 99% scaling efficiency with a 10% improvement over ZeRO-3 and a 2× improvement over ZeRO-2. This could be attributed to the fact that DDP implements AllReduce collective communication while DeepSpeed implements Reduce-Scatter and AllGather collective communication operations. The performance of the NCCL backend is highly optimized for AllReduce in comparison to AllGather. This highlights an opportunity to explore further optimizations for the DeepSpeed implementation to scale on systems. We would also like to note that there are additional tuning knobs at the NCCL layer and in DeepSpeed, and this needs further investigation for optimal performance.

For sequence length 10,240, we used a batch size of 1 and ZeRO-3. As we increased the sequence length from 2,048 to 10,240, the memory requirements, including for activation and residuals, increased by a similar factor. The computation requirements also grew by 5×. We were able to fit only one batch for this sequence length on the GPU with the current stages employed. From Fig. 6C, at 512 GPUs, for the 25M case, we observed a 50% improvement on Selene (64 nodes) over Polaris (128 nodes). For the 250M case, we observed only an 11% improvement for Selene over Polaris. As the model size increased for this sequence length, we were bottlenecked primarily by the memory subsystem performance and the overheads associated with staging residuals and parameters between the GPU and host. Additional staging optimization, model and activation partitioning will need to be explored.

### Compute Performance

We discuss the overall compute performance of the GenSLM model as we scaled the model size from 25M to 25B on the Polaris system for our production science runs. Table 3 illustrates the measured GPU performance obtained using the DeepSpeed profiler for smaller-scale runs. We next take the efficiency of the runs as we weak-scaled to larger nodes and GPU counts for the sustained PFLOPS. We would like to note that we account for the entire end-to-end application run, including data processing and checkpointing. The number of GPUs for our production science runs on Polaris was chosen based on system availability, and the number of steps run was chosen to achieve an appropriate loss scale. We observed that as we scaled the model size, the overall computational flops per step increased given the increase in model complexity. For the 25B model, we achieved a sustained performance of 44.79 PFLOPS in mixed precision (MP). We achieved a peak performance of 212.55 PFLOPS(MP) measured by accounting for the highest FLOPS consumed by a single layer in our network. For our production science runs, we used an aggregate of **1.63 Zettaflops**, and our 25B model used 1.48 Zettaflops to train on 1,024 GPUs for 2200 steps. For our scaling runs on Selene with the 25B model, we scale to 512 nodes with 4096 GPUs. We achieve a sustained performance of 121.26 PFLOPS(MP) and a peak performance of 850.21 PFLOPS(MP). The final model performance is described in Table 4.

**Table 3:** Compute performance of the production runs in mixed precision (MP) for different model sizes with a sequence length of 2,048. This includes the I/O, computations needed for forward pass, backward pass and weight updates, nnrl communication.

| Model Size | Tflops/step(MP) | sec/step | TFLOPS/GPU(MP) | # GPUs | Num steps | Sustained PFLOPS(MP) | Total Zflops(MP) |
| --- | --- | --- | --- | --- | --- | --- | --- |
| 25M | 13.2 | 0.9 | 14.7 | 512 | 3500 | 3.49 | 0.02 |
| 250M | 58.7 | 3.4 | 17.3 | 512 | 1800 | 7.59 | 0.05 |
| 2.5B | 135.3 | 4.5 | 30.3 | 256 | 2250 | 5.83 | 0.08 |
| 25B | 654.9 | 14.9 | 43.7 | 1024 | 2200 | 44.79 | 1.48 |

**Table 4:** Final loss and perplexity values achieved by the GenSLM Foundation (F) (2,048 tokens) and SARS-CoV-2 (S) (10,240 tokens) models. Reported values for S models are trained on the first year of SARS-CoV-2 genomes. Perplexity is computed by taking the exponential of the loss and can be interpreted as the number of guesses needed for the model to correctly fill a masked token.

| Metric | 25M F | 250M F | 2.5B F | 25B F | 25M S | 250M S |
| --- | --- | --- | --- | --- | --- | --- |
| Loss | 0.57 | 0.46 | 0.30 | 0.70 | 0.015 | 0.011 |
| Perplexity | 1.78 | 1.59 | 1.34 | 2.01 | 1.02 | 1.01 |

### Cerebras Wafer-Scale Cluster Scalability

We measured the throughput and training time of the Wafer-Scale Cluster for GenSLM-123M and GenSLM-1.3B with a sequence length of 10,240 codon tokens (Table 1). The batch size per CS-2 for each model was chosen based on empirical experiences with models of similar sizes and was kept constant when scaled up to multiple CS-2s.

Table 5 shows average samples per second training GenSLM-123M and GenSLM-1.3B for 200 steps using one, two, and four CS-2s. Regardless of the model configurations, we observed linear weak scaling when using up to four CS-2s.

**Table 5:**
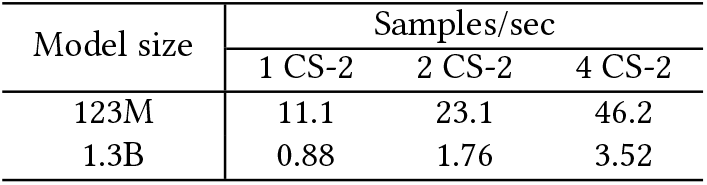
Cerebras Wafer-Scale Cluster throughput training GenSLMs on a sequence length of 10,240 tokens.

| Model size | Samples/sec |  |  |
| --- | --- | --- | --- |
|  | 1 CS-2 | 2 CS-2 | 4 CS-2 |
| 123M | 11.1 | 23.1 | 46.2 |
| 1.3B | 0.88 | 1.76 | 3.52 |

We trained GenSLM-123M and GenSLM-1.3B from scratch using learned positional embeddings. Table 6 shows training time and a total number of training samples used to achieve validation accuracy >96% and perplexity <1.03 using one CS-2 and with four CS-2s. Validation measurements were taken from checkpoints every 500 steps. For GenSLMs of the same size, fewer training steps were required to achieve comparable validation results when the global batch size was increased in a four-CS-2 Wafer-Scale Cluster with data parallelism. The reduced number of training steps plus linear weak scaling led to a reduction of at least a third of the training time when using four CS-2s versus one. All GenSLM training with full genomes on CS-2s converged within 12 hours. GenSLM-1.3B requires fewer training samples than the smaller GenSLM-123M to achieve comparable validation metrics, following the sample efficiency observation in neural language model scaling laws (Kaplan et al., 2020). We note that further hyperparameter tuning is required to 1) optimize the throughput on the Wafer-Scale Cluster, 2) draw firmer conclusions on the impact of model size on model quality.

**Table 6:** Metrics of GenSLMs trained from scratch on a sequence length of 10,240 using Cerebras Wafer-Scale Cluster.

|  | GenSLM-123M |  | GenSLM-1.3B |  |
| --- | --- | --- | --- | --- |
|  | 1 CS-2 | 4 CS-2 | 1 CS-2 | 4 CS-2 |
| Training steps | 5,000 | 3,000 | 4,500 | 3,000 |
| Training samples | 165,000 | 396,000 | 49,500 | 132,000 |
| Time to train (h) | 4.1 | 2.4 | 15.6 | 10.4 |
| Validation accuracy | 0.9615 | 0.9625 | 0.9622 | 0.9947 |
| Validation perplexity | 1.0310 | 1.0290 | 1.0310 | 1.0255 |

### Workflow Performance

We measured the utilization of the sequence generation workflow on 224 nodes of Polaris by counting the number of workers actively serving a request as a function of runtime. As shown in Fig. 7, we achieve 97.0% utilization over the 5.5-hour duration of the workflow. Persisting the GenSLMs in GPU memory between requests generated 3.85 sequences per second, whereas without model caching we estimate the workflow would have only generated 1.98 sequences per second by extrapolating the mean cold start time across the number of workers. This achieves 1.9× faster time to solution for generating synthetic sequences with notable properties, allowing for rapid analysis at time scales not previously feasible.

**Figure 7:**
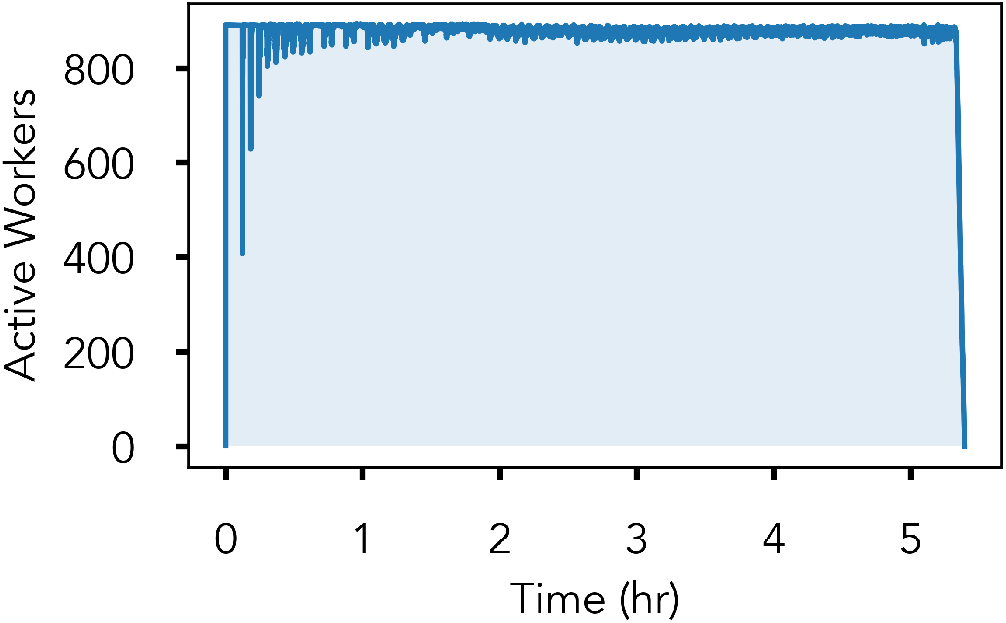
Workflow utilization measured by the number of active workers (applications actively serving requests) as a function of workflow runtime measured on 224 nodes of Polaris (896 A100 GPUs). The warm-able application design realizes 97% utilization, enabling 1.9X more sequences to be generated compared to a cold start baseline.

## 8 IMPLICATIONS

In this paper, we presented GenSLMs, one of the first LLMs trained on nucleotide sequences, particularly at the genome scale, and demonstrated its performance in modeling evolutionary dynamics of SARS-CoV-2. Our approach overcomes key challenges related to training LLMs for biological data, specifically with respect to longer sequence lengths and building biologically meaningful latent spaces which can then be used for a variety of downstream prediction tasks. GenSLM is a foundation model for biological sequence data and opens up avenues for building hierarchical AI models for several biological applications, including protein annotation workflows, metagenome reconstruction, protein engineering, and biological pathway design. We scaled the training of GenSLM for sequence length up to 10240 tokens and 25B parameters on GPU-based supercomputers. We scaled to 4,096 GPUs and utilized over 1.63 Zettaflops for science runs. We identified scaling avenues to be pursued in order to tackle larger models and sequence lengths needs for science. We demonstrated the efficacy of the Cerebras Wafer-Scale Cluster, an AI accelerator, to scale the training of GenSLM with high user-productivity and achieved linear scaling for 10,240 tokens and models up to 1.3B parameters.

We also note that the information contained within nucleotide sequences represents a much richer vocabulary compared to PLMs alone. Thus, the learned representation lets us capture a much larger repertoire of biological properties that are perhaps diminished while using PLMs, and enables a more faithful generation process that captures the intrinsic organization of the SARS-CoV-2 sequences. Further, the attention mechanism also reveals co-evolutionary patterns at the whole-genome scale that requires future investigation to fully understand how these long-range interactions may influence our ability to inform epitope modeling, immune escape, antibody design, and even vaccine design strategies. We however note that there is a need to rigorously compare PLMs with GenSLM-like approaches. It remains to be seen if the GenSLM model does possess richer representative power and if so how it can be further used. Note that we have also not been able to address the aspects of noise and bias in the data – similar to natural language models where the models demonstrated extreme bias, there needs to be rigorous analyses of GenSLMs generative capabilities. We welcome the community to drive the development of suitable test harnesses for rigorously evaluating GenSLM-like models.

A straightforward extension to our work would include the integration of GenSLMs with protein structure prediction workflows such as AlphaFold (Jumper et al., 2021)/OpenFold^4^ and faster protein folding methods (Lin et al., 2022) to model both immune escape and fitness, which determine the ability of the virus to adapt to its host (human) (Beguir et al., 2022). Further, incorporating experimental and biophysical data into our workflow from antibody binding assays, molecular docking, and other quantitative metrics can also guide the training regimes for these models such that the generative process can be constrained to focus on potential future variants of concern.

## ACM Reference Format

Maxim Zvyagin^1 †^, Alexander Brace^1,2†^, Kyle Hippe^1†^, Yuntian Deng^3,4†^, Bin Zhang^5^, Cindy Orozco Bohorquez^5^, Austin Clyde^1,2^, Bharat Kale^6^, Danilo Perez-Rivera^1,7^, Heng Ma^1^, Carla M. Mann^1,2^, Michael Irvin^1^, J. Gregory Pauloski^2^, Logan Ward^1^, Valerie Hayot-Sasson^1,2^, Murali Emani^1^, Sam Foreman^1^, Zhen Xie^1^, Diangen Lin^1,2^, Maulik Shukla^1,2^, Weili Nie^3^, Josh Romero^3^, Christian Dallago^3,9^, Arash Vahdat^3^, Chaowei Xiao^8,3^, Thomas Gibbs^3^, Ian Foster^1,2^, James J. Davis^1,2^, Michael E. Papka^1,10^, Thomas Brettin^1^, Rick Stevens^1,2^, Anima Anandkumar^3,11^*, Venkatram Vishwanath^1^*, Arvind Ramanathan^1^*. 2020. GenSLMs: Genome-scale language models reveal SARS-CoV-2 evolutionary dynamics. In *Supercomputing ‘22: International Conference for High Performance Computing, Networking, Storage, and Anal-ysis*. ACM, New York, NY, USA, 14 pages. https://doi.org/finalDOI

## ACKNOWLEDGMENTS

We thank the Argonne Leadership Computing Facility (ALCF) supported by the DOE under DE-AC02-06CH11357 and the National Energy Research Scientific Computing Center (NERSC) at Lawrence Berkeley National Laboratory supported by the DOE under Contract No. DE-AC02-05CH11231. We thank Bill Allcock, Silvio Rizzi and ALCF, Wahid Bhimji and NERSC for their timely help in enabling us to run these jobs at scale. We also thank Defne Gorgun, Lorenzo Casalino and Rommie Amaro for stimulating discussions. This research was supported by the Exascale Computing Project (17-SC-20-SC), a collaborative effort of the US DOE Office of Science and the National Nuclear Security Administration, the National Institute of Allergy and Infectious Diseases, National Institutes of Health Award Number P01AI165077 (AR), the National Science Foundation Award Number 2117896 and supported by the DOE through the National Virtual Biotechnology Laboratory, a consortium of DOE national laboratories focused on response to COVID-19, with funding from the Coronavirus CARES Act.

## Footnotes

1 https://www.gisaid.org

2 https://github.com/ramanathanlab/genslm

3 https://www.bv-brc.org

4 http://openfold.io

